# A shared molecular target but distinct genetic routes to diamide resistance in *Helicoverpa armigera* (Lepidoptera: Noctuidae)

**DOI:** 10.64898/2026.10.13.758793

**Authors:** Douglas Amado, Anne Caroline Santos Küll, Fábio Miguel Führ, Victor Flores, Erika Pinheiro de Castro, Alberto Soares Corrêa, Chris D. Jiggins, Celso Omoto

## Abstract

Diamide insecticides target insect ryanodine receptors (*RyR*s) and are widely used against lepidopteran pests, yet it remains unclear whether compounds sharing this molecular target evolve resistance through the same genetic routes. Here, we integrated *RyR* sequencing, structural modelling and population genomics to investigate resistance to chlorantraniliprole and flubendiamide in the global agricultural pest *Helicoverpa armigera*. We identified G4946E together with F4947I, a previously undescribed resistance-associated *RyR* substitution, in a chlorantraniliprole-resistant strain. Structural modelling indicated that F4947I may modify G4946E’s effects on the diamide-binding pocket, while field and population-genomic evidence independently implicated the *RyR*-containing chromosome 28 region in chlorantraniliprole resistance. By contrast, high-level flubendiamide resistance occurred without amino acid substitutions in the *RyR* region examined, and field survival was predominantly associated with chromosome 2, independently recapitulating a previously identified resistance locus. Together, our findings reveal that resistance to insecticides sharing the same molecular target can evolve through distinct but potentially overlapping genetic routes, highlighting the limitations of predicting cross-resistance from target-site variation alone and the need for broader genomic approaches to resistance surveillance.

**Graphical Abstract:** 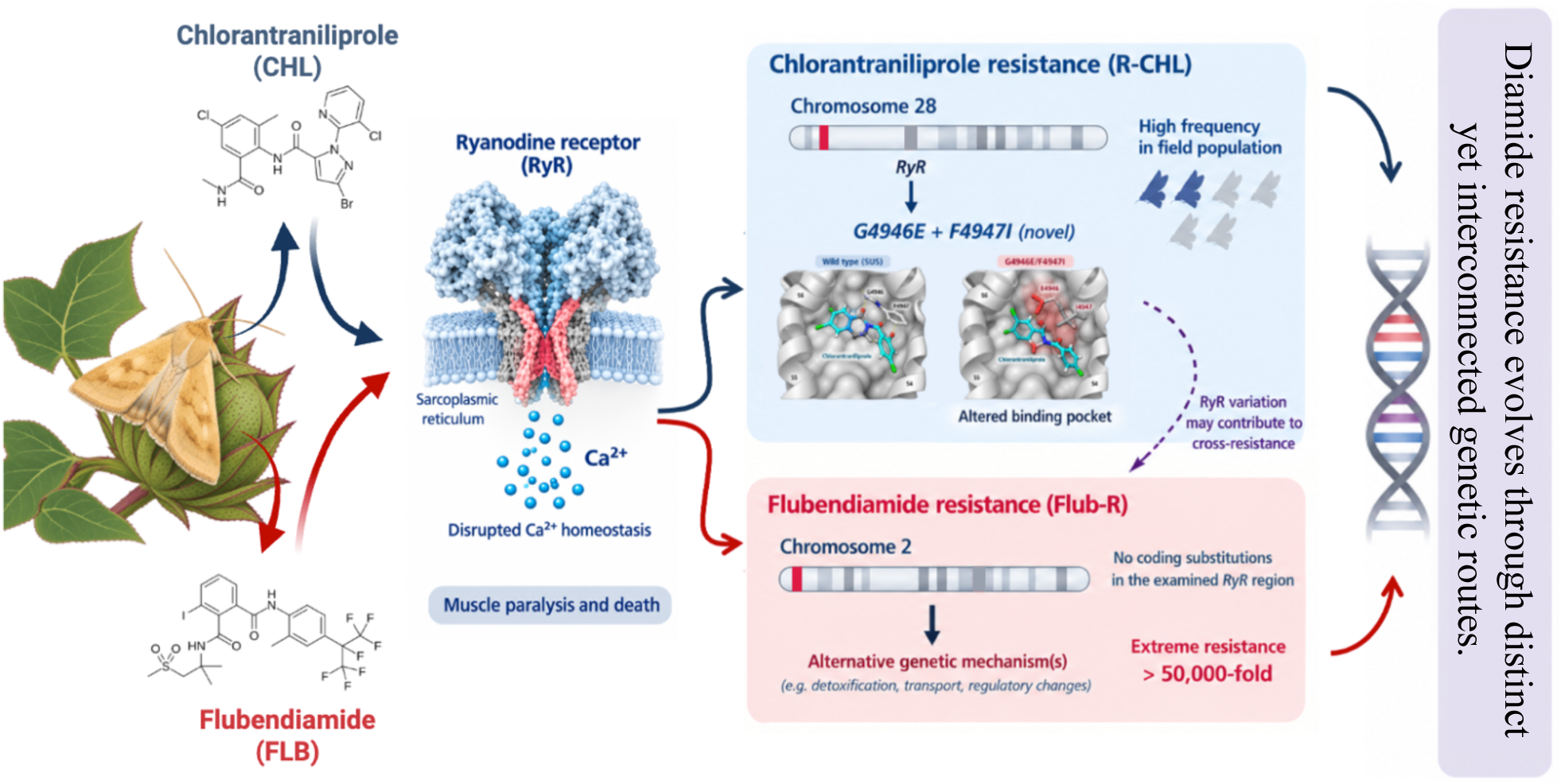

## Introduction

The intensive use of insecticides in agriculture is driven, in part, by the rapid evolution of insect resistance, a process that reduces control efficacy, increases environmental burdens, promotes productivity losses, and ultimately threatens food security (Bakker et al. 2020; Ma et al. 2021; Wan et al. 2025). This problem is especially acute for the key pest *Helicoverpa armigera* (Hübner, 1805) (Lepidoptera: Noctuidae), a major contributor to yield losses across intensive cropping systems in Asia, Oceania and the Americas (Riaz et al. 2021; Jin et al. 2023). Its global success reflects phenotypic plasticity, a broad host range and, in the Americas, bidirectional interspecific gene flow with its sister species *Helicoverpa zea* (Boddie, 1850) (Lepidoptera: Noctuidae) (Anderson et al. 2018; Cordeiro et al. 2020; North et al. 2025), which may facilitate the introgression of resistance genes into species where they were previously absent, thereby accelerating their spread. Currently, managing agricultural pest species relies on a relatively limited range of insecticides, as discovering and developing new insecticidal chemistries has become increasingly difficult (Sparks and Bryant 2022; Sparks 2025). Among the classes that remain widely effective, diamides are central to chemical control. Any erosion of their performance, therefore, has immediate consequences for production systems and the long-term sustainability of integrated pest management.

Diamide insecticides selectively activate insect ryanodine receptors (*RyR*s), disrupting intracellular Ca²⁺ homeostasis through sustained release of calcium from intracellular stores. The resulting depletion of Ca²⁺ reserves causes rapid muscle dysfunction, feeding cessation, paralysis, and ultimately death (Cordova et al. 2006; Ebbinghaus-Kintscher et al. 2006; Nauen 2006; Lahm et al. 2009). *RyR*s are large tetrameric Ca²⁺-release channels with substantial regulatory complexity and constitute the primary molecular target of this insecticide class (Cordova et al. 2006; Ebbinghaus-Kintscher et al. 2006; Jeanguenat 2013; Ma et al. 2020; Richardson et al. 2020; Du and Fu 2023). Resistance to diamides can arise through two broad routes: (1) target-site mechanisms, particularly mutations in *RyR*, and (2) non-target-site mechanisms, including enhanced metabolic detoxification (e.g., cytochrome P450 monooxygenases, esterases and glutathione S-transferases), altered penetration and xenobiotic transport by ATP-binding cassette (ABC) transporters (Han et al. 2023; Han et al. 2024; Amado et al. 2025). In Lepidoptera, *RyR* mutations represent one of the best-characterised mechanisms of diamide resistance and are frequently concentrated within transmembrane regions, including the pseudo-voltage-sensor domain (*pVSD*), where substitutions can alter the structural and functional properties of the diamide-binding region (Ma et al. 2020; Lin et al. 2024).

*RyR* variants as G4946E, I4790K have been associated with reduced susceptibility to diamides across multiple species (Troczka et al. 2012; Tao et al. 2013; Nauen and Steinbach 2016; Roditakis et al. 2017; Boaventura et al. 2020; Jouraku et al. 2020a; Richardson et al. 2020; Zuo et al. 2020; Dunn et al. 2022; “Sam” Dunn et al. 2025). Notably, the phthalic diamide flubendiamide often exhibits substantially higher resistance ratios than anthranilic diamides such as chlorantraniliprole (Troczka et al. 2012; Steinbach et al. 2015a; Troczka et al. 2015), indicating that resistance phenotypes can differ markedly even among insecticides acting on the same molecular target. Additionally, combinations of *RyR* mutations can increase resistance levels and/or confer cross-resistance (Guo et al. 2014; Huang et al. 2021). Recent structural studies have begun to reveal how individual and combined substitutions can alter the architecture and functional properties of the diamide-binding region (Lin et al. 2024). However, it remains unclear whether resistance to different diamides is predominantly governed by a common genetic architecture centred on *RyR*, or whether individual compounds can favour partially distinct evolutionary routes to resistance.

Population genomic and genotype–phenotype association approaches provide a powerful framework for resolving the genetic architecture of insecticide resistance beyond individual candidate mechanisms (Cosme et al. 2022; Lucas et al. 2023; Amado et al. 2025). This is particularly important in field populations, where resistance phenotypes may reflect standing genetic variation and the combined effects of target-site and non-target-site mechanisms evolving under heterogeneous selection pressures (Gadji et al. 2024). By linking genome-wide variation directly to differences in insecticide survival across natural populations, these approaches can recover known resistance loci while also identifying additional genomic regions that established mechanisms alone would not necessarily predict (Lucas et al. 2023; Amado et al. 2025). Such analyses are therefore especially valuable for insecticides sharing the same molecular target, as they allow testing whether resistance to different compounds is associated with a common genetic architecture or with distinct genomic routes (Zuo et al. 2017; Jouraku et al. 2020b; Jiang et al. 2021). Examining naturally occurring variation in field populations can therefore provide a more comprehensive view of the genetic diversity underlying resistance and the mechanisms potentially contributing to its emergence, persistence, and spread (Clarkson et al. 2021).

These questions are particularly relevant in *H. armigera*, where our previous genetic analyses identified a major genomic region on chromosome 2 associated with flubendiamide resistance, even though *RyR* is located on chromosome 28. This raises the possibility that high-level resistance can evolve through mechanisms outside the canonical target-site pathway (Amado et al. 2025). While the flubendiamide-resistant strain showed no cross-resistance to chlorantraniliprole (Abbade-Neto et al. 2022), a recently characterized chlorantraniliprole-resistant strain displayed high cross-resistance to flubendiamide (Führ et al. 2026). Resolving how *RyR* variation and alternative genomic mechanisms contribute to resistance across distinct diamides is therefore essential for understanding the evolution of cross-resistance and for determining whether resistance to compounds sharing the same molecular target can be reliably predicted from target-site variation alone.

Thus, in this study, we integrated targeted *RyR* sequencing, structural modelling and population-genomic analyses to investigate the genetic architecture of diamide resistance in *H. armigera*. Specifically, we first characterised variation within the transmembrane region of *RyR* in strains exhibiting contrasting resistance phenotypes and evaluated the potential structural consequences of resistance-associated amino acid substitutions for chlorantraniliprole binding. We then examined the distribution of these *RyR* variants in field-derived populations and integrated genome-wide differentiation and genotype–phenotype association analyses to identify genomic regions associated with survival to chlorantraniliprole and flubendiamide. By comparing evidence from laboratory-selected strains with genomic variation in field populations, we aimed to determine the extent to which resistance to these two diamides reflects shared target-site mechanisms or alternative, compound-specific genetic routes. Resolving this distinction is critical for understanding the evolution of diamide resistance and for developing molecular surveillance strategies that can detect the diversity of resistance mechanisms present in field populations.

## Results

### Two mutations found in *RyR* associated with resistance

To identify non-synonymous variants associated with diamide resistance, we sequenced overlapping PCR amplicons covering a region of the *RyR* gene from different laboratory strains of *H. armigera*. The resulting consensus spans 517 amino acids and encompasses transmembrane domains II-VI. No amino acid substitutions were detected in the SUS strain (Fig. 1). Similarly, no amino acid-altering variants were detected in the Flub-R strain. By contrast, the R-CHL strain harboured two adjacent amino-acid substitutions: G4924E and F4925I, corresponding to G4946E and F4947I in *P. xylostella* numbering, respectively (Fig. 1). Thus, the high-level flubendiamide resistance previously reported in the Flub-R strain is not explained by sequence changes within this *RyR* segment. Paradoxically, R-CHL, which carries two adjacent amino-acid substitutions in the same segment, also exhibited a flubendiamide resistance ratio exceeding 23,000-fold, indicating that flubendiamide resistance cannot be straightforwardly inferred from coding variation in this region alone.

**Figure 1.**
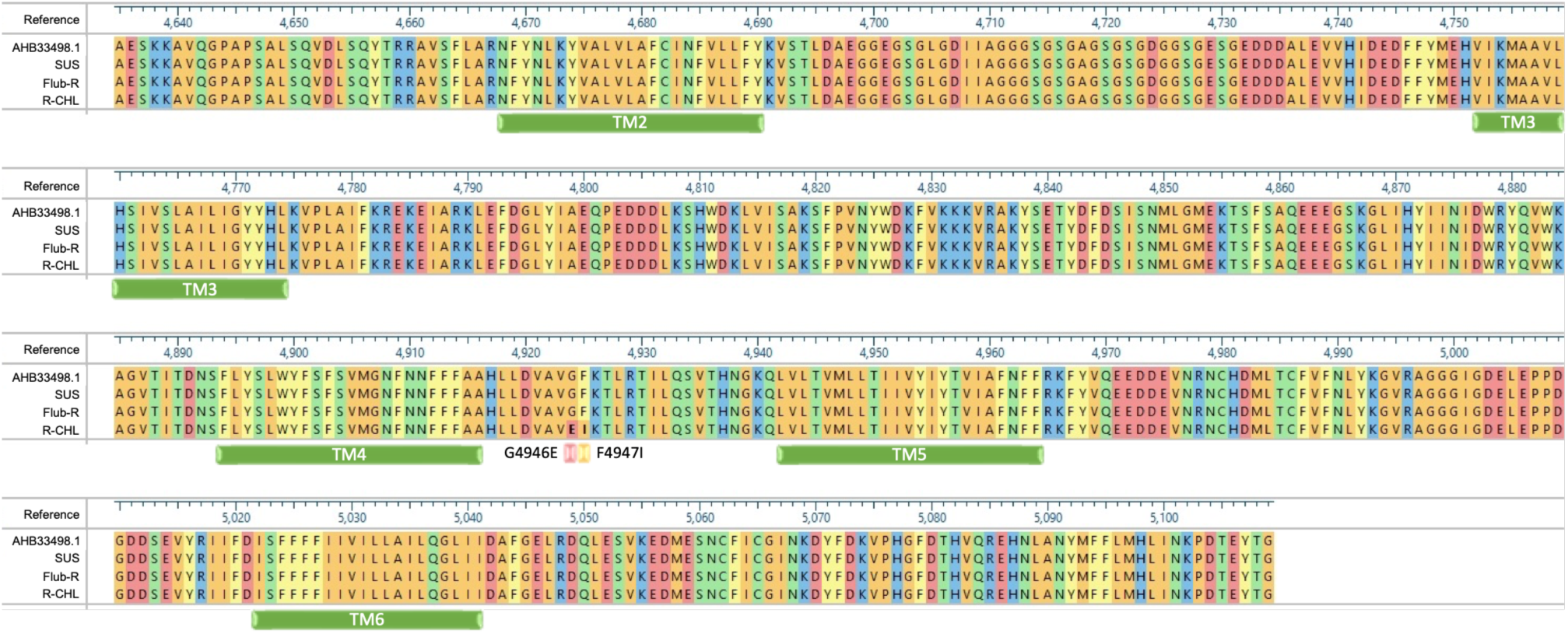
Multiple sequence alignment of the *RyR* segment spanning transmembrane domains II-VI identifying two adjacent substitutions (G4946E and F4947I) in the R-CHL strain. Multiple sequence alignment of a partial *H. armigera* ryanodine receptor (*RyR*) segment (≈500 amino acids) encompassing transmembrane domains II-VI. The alignment was generated using the *H. armigera RyR* sequence (GenBank: AHB33498.1) and compared against the homologous region of *Plutella xylostella RyR* (GenBank: AET09964.1). Two adjacent amino acid substitutions were detected in the chlorantraniliprole-resistant strain (R-CHL), G4946E and F4947I. These substitutions were absent in the susceptible (SUS) and flubendiamide-resistant (Flub-R) strains, previously shown by Abbade-Neto et al. (2022). The alignment indicates substitution positions.

### Mutations associated with resistance disrupt the *RyR* binding to chlorantraniliprole

We performed structural modelling using AlphaFold3 and benchmarked it against the *RyR* cryo-EM system reported by (Lin et al. 2024). The experimentally resolved tetraniliprole-*RyR* complex (Fig. 2A) and the same *RyR* sequence from Lin et al. *RyR* sequence modelled with chlorantraniliprole (Fig. 2B) was used as a positive control, providing an internal consistency check that the predicted ligand pose localises to the diamide-binding pocket defined by the cryo-EM template. Within this framework, the models predicted contrasting effects of the two substitutions on the chlorantraniliprole-binding site (Fig. 2). Relative to the susceptible model, G4946E produced the most pronounced structural impact, reshaping the *RyR* binding pocket in a manner consistent with reduced ligand accommodation of chlorantraniliprole (Fig. 2D). By contrast, F4947I had a limited effect, largely preserving pocket geometry and the predicted pose of chlorantraniliprole with respect to key binding residues (Fig. 2E). Notably, in the G4946E/F4947I double mutant from the resistant R-CHL strain, the model indicated occlusion of the binding site and a ligand pose incompatible with stable binding, suggesting loss of pocket accessibility to chlorantraniliprole (Fig. 2F).

**Figure 2.**
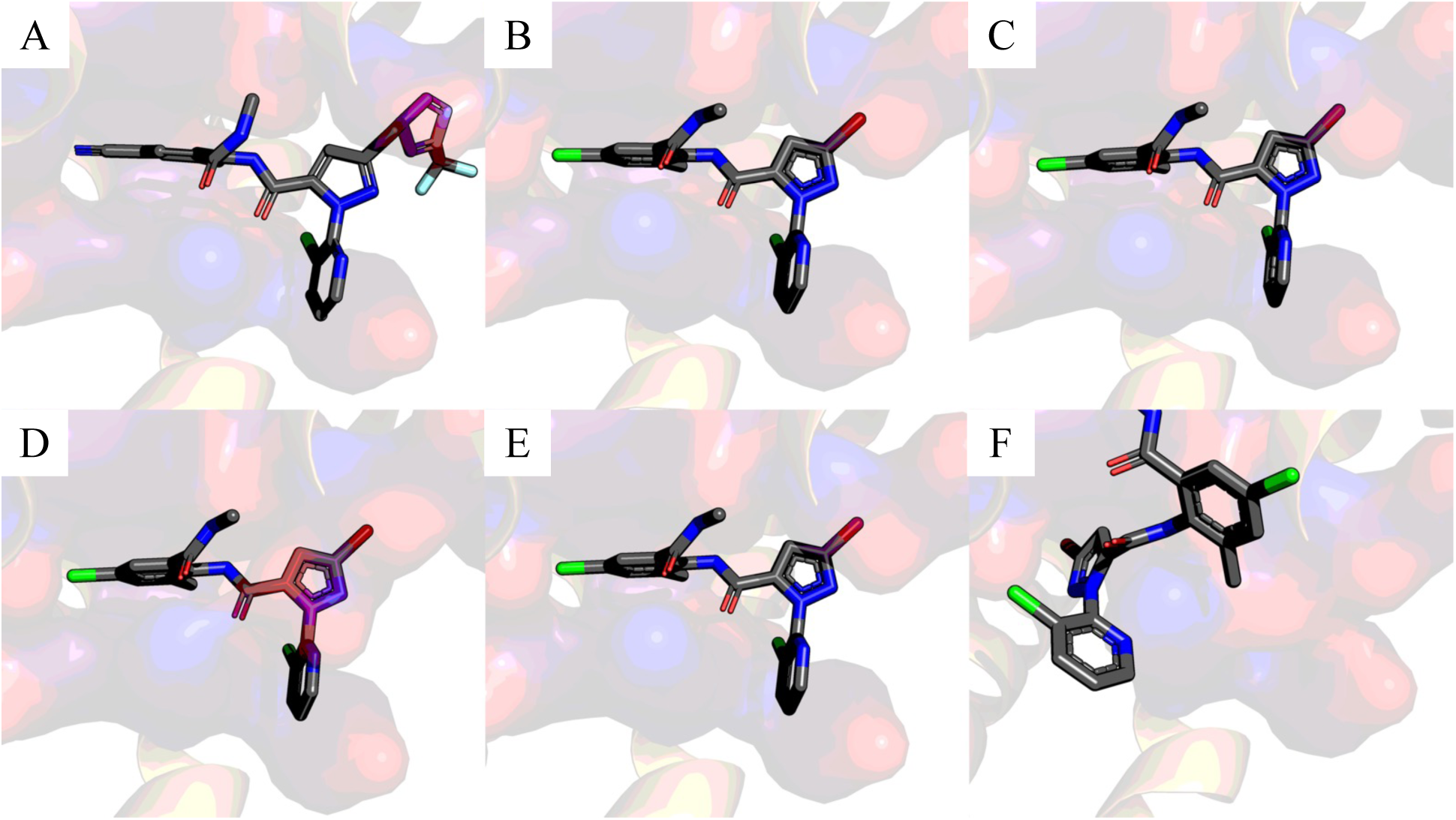
Structural models of the ryanodine receptor (*RyR*) binding pocket with the insecticide chlorantraniliprole, which predicts disruption of the binding in G4946E and the double mutant, but not in F4947I. Cryo-EM *RyR* model from Lin et al. (2024) in with tetraniliprole (A) and (B) with chlorantraniliprole. C) Structural model of *RyR* without the mutations G4946E and F4947I, which is found in the *H. armigera* susceptible strain (SUS), with chlorantraniliprole as a ligand. D) G4946E single mutant model (derived from the R-CHL background) docked with chlorantraniliprole. E) F4947I single mutant model (derived from the R-CHL background) docked with chlorantraniliprole. F) R-CHL resistant-strain *RyR* double mutant (G4946E/F4947I) docked with chlorantraniliprole. In all panels, the ligand is shown in grey and predicted pocket residues are shown in purple.

### Field Population Screening

Of the 221 individuals from the 12 populations tested, we identified 201 homozygous susceptible, 8 homozygous resistant, and 12 heterozygous individuals (Table 1). We first identified the resistance allele in Bahia in the 2017 crop season and then in Mato Grosso in the 2018 crop season (Table 1). Screening focused on populations from these two Brazilian states, reflecting the geographical origins of the laboratory-selected resistant strains: R-CHL, derived from Mato Grosso, and Flub-R, derived from Bahia.

**Table 1.** Field populations of *H. armigera* screened across crop seasons for susceptibility monitoring. Field populations collected across crop seasons (2015-2021) and locations (latitude/longitude) and assayed for susceptibility monitoring. The table reports survival (%) and genotype counts for susceptible homozygotes (Sus Hom.), resistant homozygotes (Res Hom.) and heterozygotes (Het.). Totals are shown in the final column.

| Pop. | Crop Season | Latitude (S) | Longitude (W) | Survival % | Sus Hom. | Res Hom. | Het. | Total |
| --- | --- | --- | --- | --- | --- | --- | --- | --- |
| BA01 | 2015 | 12°42'02" | 48°02'17" | 2.50 | 16 | 0 | 0 | 16 |
| BA02 | 2016 | 12°05'58" | 55°10'08" | 7.50 | 12 | 0 | 0 | 12 |
| BA03 | 2016 | 12°05'58" | 52°33'04" | 5.95 | 16 | 0 | 0 | 16 |
| BA04 | 2016 | 11°46'40" | 45°35'39" | 4.24 | 19 | 0 | 0 | 19 |
| BA05 | 2017 | 11°51'23" | 47°45'38" | 25.86 | 17 | 1 | 0 | 18 |
| BA06 | 2019 | 12°45'00" | 46°33'16" | 21.67 | 16 | 1 | 6 | 23 |
| MT01 | 2016 | 15°33'32" | 54°46'46" | 1.25 | 22 | 0 | 0 | 22 |
| MT02 | 2017 | 15°24'41" | 47°17'53" | 3.33 | 11 | 0 | 0 | 11 |
| MT03 | 2017 | 16°21'09" | 52°52'57" | 10.49 | 24 | 0 | 0 | 24 |
| MT04 | 2018 | 17°04'24" | 58°39'26" | 8.75 | 18 | 0 | 0 | 18 |
| MT05 | 2018 | 13°32'42" | 45°43'43" | 4.16 | 19 | 0 | 1 | 20 |
| MT06 | 2021 | 17°09'36" | 47°54'04" | 30.83 | 11 | 6 | 5 | 22 |
| **Total** | - | - | - | - | **201** | **8** | **12** | **221** |

The correlation analysis between the resistance allele frequency and the survival percentage across the 12 populations yielded a strong positive correlation coefficient of 0.82 [t(10) = 4.56, p = 0.001] (Fig. S1A). A strong positive correlation (correlation coefficient= 0.86) was also observed between resistance allele frequency and crop seasons [t(10) = 5.36, p-value < 0.001] (Fig. S1B). Together, these analyses suggest that the allele conferring resistance to the insecticide chlorantraniliprole emerged and increased in frequency in field populations of *H. armigera* in the 2018/2019 crop season.

Using ROC analysis, the molecular marker yielded an AUC of 1 and a p-value of < 0.001. The optimal threshold was 0.0403, yielding perfect sensitivity and specificity (both 1). These results suggest that, within the analysed dataset, the marker ideally discriminates between individuals resistant and those susceptible to chlorantraniliprole.

### F_ST_ scan across field populations

We performed an *F_ST_* scan using 189 individuals and, after filtering, included 1,909,879 SNPs in the analysis. Based on our classification, Population A (survival rates ranging from 1% to 10%) comprised 116 individuals, and Population B (survival rates of 11% or higher) comprised 73 individuals for chlorantraniliprole. For flubendiamide, Population A (survival rates ranging from 1% to 14%) comprised 75 individuals, and Population B (survival rates of 20% or higher) comprised 114 individuals (Table S2).

Across most of the genome, *F_ST_* values were low, indicating broadly modest differentiation among sampled field populations. In contrast, we observed two pronounced outlier regions on chromosomes 2 and 28 (Fig. 3).

**Figure 3.**
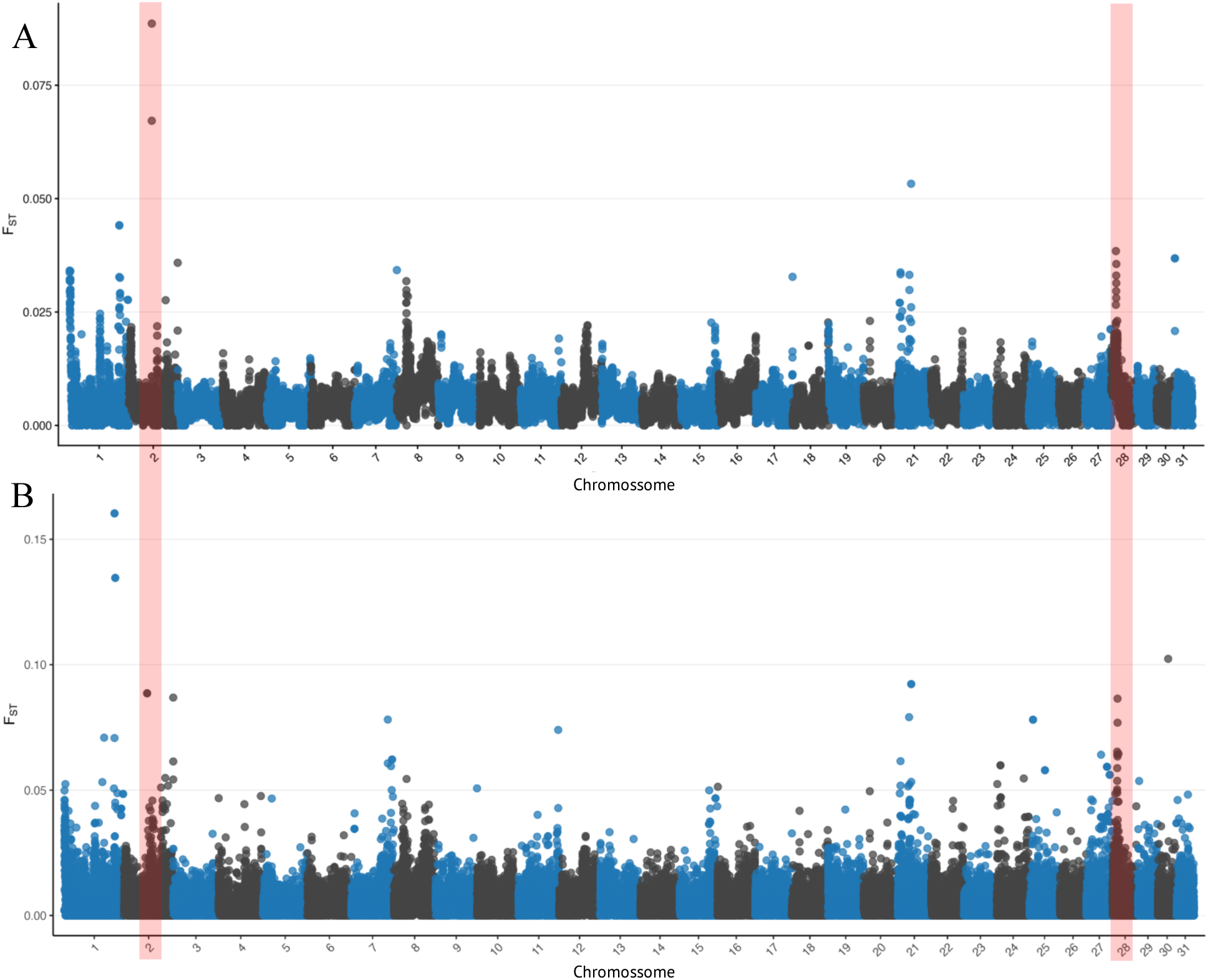
F_ST_ scan across field populations collected from 2014-2021 crop seasons. Genome-wide F_ST_ scan calculated across 189 individuals included in the two-population dataset, with stratification by phenotype. A) Window of 50 kb and Step of 20 kb; B) Window of 10 kb and Step of 4 kb. Each point represents the estimated *F_ST_* for a variant along the genome, plotted by chromosomal position. The red boxes highlight genomic regions that remained significant after reducing the window size from 50 kb to 10 kb, indicating that these signals are robust and unlikely to be false positives.

On chromosome 2, a sharp, narrow *F_ST_* peak rises well above the genomic background, reaching the highest differentiation levels observed in the scan (Fig. 3). The steep, localised profile is consistent with a restricted genomic interval exhibiting markedly different allele frequencies among populations. Because the analysis does not condition on survival phenotype, this signal reflects population-level differentiation at this locus, potentially driven by geographically structured selection or historical/demographic processes, rather than a direct case-control contrast.

On chromosome 28, a second prominent peak is evident, with elevated *F_ST_* concentrated in a discrete region and standing out against the low genome-wide baseline (Fig. 3). The magnitude and localisation of this peak are consistent with large among-population differences in allele frequencies at, or near, a major resistance-associated locus. Given the established importance of chromosome 28 for diamide resistance (including the *RyR* region), this differentiation signal is compatible with spatially heterogeneous selection on resistance variants across Brazilian field populations over the 2014-2021 sampling period. Together, the chromosome 2 and chromosome 28 peaks indicate that key resistance-relevant genomic regions are also among the most differentiated across field populations.

### Genotype-phenotype associations with diamide survival from field populations

We performed Genome−Environment Association (GEA) using whole-genome sequencing (WGS) data from 189 *H. armigera* individuals sampled across 28 field populations collected during the 2014-2021 crop seasons, the same populations used in the *F_ST_* analysis. For each diamide insecticide (chlorantraniliprole and flubendiamide), population-level survival percentages at the corresponding diagnostic concentrations were used as phenotypic covariates (Table S2). These phenotypic data comprised previously published susceptibility-monitoring results from the 2014-2018 crop seasons (Pereira et al. 2020) and newly generated data from the 2019-2021 crop seasons, reported here for the first time. Following quality filtering, the final dataset comprised 189 individuals and approximately 1.9 million SNPs for the association analyses of both insecticides.

BayPass analyses identified SNPs whose allele frequencies were associated with variation in survival to chlorantraniliprole and flubendiamide among field populations (Fig. S3). For chlorantraniliprole, two SNPs with the strongest association exceeded the Bayes factor threshold of 20 dB, located on chromosomes 14 and 28 (Fig. S3A). In the 300 kb window around the marker on chromosome 14, we identified genes associated with diamide insecticide detoxification, such as peroxidase (LOC110380531) and glutaredoxin-3 (LOC110380771) (Table S3). The significant SNP on chromosome 28 was located within the genomic region containing the ryanodine receptor (*RyR*) gene, the molecular target of diamide insecticides.

In contrast, flubendiamide survival was associated with a broader set of genomic signals, with SNPs exceeding the 20-dB threshold distributed across multiple chromosomes (Fig. S3B). We detected the strongest association on chromosome 2. We also detected a strong association on chromosome 28, where the *RyR* gene is located. Additional above-threshold associations were distributed across several other chromosomes, indicating that variation in flubendiamide survival is associated with multiple genomic regions.

Collectively, the BayPass analyses revealed both shared and insecticide-specific genomic associations. Chromosome 28 was associated with survival to both chlorantraniliprole and flubendiamide and encompasses the locus of the *RyR* substitutions G4946E and F4947I, supporting an association between this locus and survival in field populations following exposure to chlorantraniliprole and flubendiamide (Fig. S3A & S3B). The prominent chromosome 2 signal was particularly evident for flubendiamide and recapitulated the genomic region previously associated with resistance to this insecticide. Together, these field-based results indicate that distinct resistance mechanisms co-occurred, acting in different genomic regions and dispersing across Brazil through time.

## Discussion

Our results reveal that resistance to diamide insecticides in *H. armigera* is underpinned by a heterogeneous genetic architecture rather than by a single, conserved resistance mechanism across compounds (Huang et al. 2020; Jiang et al. 2021; Kim et al. 2025). For chlorantraniliprole, multiple independent lines of evidence converge on a role for target-site variation in the ryanodine receptor (*RyR*): the presence of the adjacent G4946E and F4947I substitutions in the resistant R-CHL strain; elevated genomic differentiation on chromosome 28; an association between survival and variation within the *RyR*-containing region; and structural predictions indicating disruption of the diamide-binding pocket of *RyR*. In contrast, flubendiamide resistance showed a markedly different genetic pattern: despite the absence of amino acid substitutions in the examined *RyR* transmembrane region, the Flub-R strain exhibited high-level resistance (Amado et al. 2025). Field survival was also strongly associated with chromosome 2, independently supporting a genomic region previously implicated in flubendiamide resistance (Abbade-Neto et al. 2022; Amado et al. 2025). Together, these findings indicate that resistance to insecticides sharing the same molecular target can evolve through partially distinct genetic routes: with target-site resistance contributing prominently to chlorantraniliprole resistance, whereas alternative mechanisms independent of coding variation within the examined *RyR* region appear to play a major role in flubendiamide resistance.

The target-site component identified here is consistent with the well-established role of *RyR* variation in the evolution of diamide resistance across lepidopteran pests. The mutation G4946 (using *P. xylostella* numbering) is one of the most extensively characterised *RyR* residues involved in diamide resistance, with substitutions at this position repeatedly identified in resistant lepidopteran populations and functionally validated through receptor expression, ligand-binding assays, and genome editing (Troczka et al. 2012; Steinbach et al. 2015a; Troczka et al. 2015; Jouraku et al. 2020a). Researchers identified the G4946E substitution in highly resistant field populations of *P. xylostella*, *Spodoptera exigua* (Hübner, 1808) (Lepidoptera: Noctuidae), *Chilo suppressalis* (Walker, 1863) (Lepidoptera: Crambidae), and *Tuta absoluta* (Meyrick, 1917) (Lepidoptera: Gelechiidae), with a similar reduction in *RyR* sensitivity to both anthranilic and phthalic diamides (Troczka et al. 2012; Troczka et al. 2015; Roditakis et al. 2017; Yao et al. 2017; Wang et al. 2026). Its occurrence in the chlorantraniliprole-selected R-CHL strain of *H. armigera* therefore provides compelling evidence that a resistance-associated substitution at this highly conserved position has arisen in another major lepidopteran pest.

Particularly, G4946E occurred together with the adjacent F4947I substitution in R-CHL. Whereas G4946E is a well-established diamide-resistance mutation across lepidopteran species, our study identifies F4947I as a previously undescribed *RyR* substitution associated with diamide resistance. Recent cryo-EM structures show that G4946E alters the local conformation and polarity of the *RyR*-binding cavity, reducing chlorantraniliprole affinity and shifting its binding pose (Lin et al. 2024). Consistent with these observations, our structural models predicted that G4946E markedly reshapes the binding pocket and disrupts chlorantraniliprole accommodation, an effect consistent with the introduction of a larger, negatively charged glutamate side chain at this functionally important position (Qi and Casida 2013; Jiang et al. 2021; Hadiatullah et al. 2022; Wang et al. 2026). By contrast, F4947I alone produced comparatively limited structural perturbation, largely preserving the binding-pocket geometry and predicted ligand pose. Strikingly, however, the combined G4946E/F4947I substitutions produced a substantially greater predicted perturbation, characterised by occlusion of the binding cavity and displacement of chlorantraniliprole from its predicted binding position. Although these structural predictions require functional validation, they identify F4947I as a novel resistance-associated candidate and suggest that this substitution modifies G4946E, enhancing disruption of diamide binding when both substitutions occur together.

Importantly, evidence for the involvement of *RyR* in chlorantraniliprole resistance was not restricted to the laboratory-selected R-CHL strain. Independent analyses of field populations converged on the same genomic region. Targeted genotyping detected the resistance-associated *RyR* alleles in four geographically distinct field-derived populations, showing that these variants are not restricted to the laboratory-selected R-CHL strain but also segregate in field populations. Similar geographic variation in the occurrence and frequency of resistance-associated *RyR* alleles has been documented in field populations of other lepidopteran pests, including *P. xylostella* and *S. frugiperda*, where G4946 and I4790 variants have been detected across multiple populations and, in some cases, associated with variation in diamide susceptibility (Okuma et al. 2022; Jiang et al. 2024).

At the genome-wide level, chromosome 28, which contains the *RyR* locus, harboured one of the two most pronounced regions of genetic differentiation among field populations, against a background of otherwise generally low F_ST_. Furthermore, our genotype–phenotype association analysis independently identified a SNP on chromosome 28 within the *RyR*-containing genomic region as associated with chlorantraniliprole survival. Together, targeted genotyping, population differentiation, and genotype–phenotype association provide complementary evidence linking variation in the *RyR* region to chlorantraniliprole resistance under field conditions. When considered together with the occurrence of G4946E/F4947I in R-CHL and the predicted structural consequences of these substitutions, these results support a model in which target-site variation at *RyR* represents an important component of chlorantraniliprole resistance in Brazilian populations of *H. armigera*, while not excluding contributions from additional resistance loci.

Despite convergent evidence implicating *RyR*, the strongest genotype–phenotype association with chlorantraniliprole survival was detected on chromosome 14, suggesting that target-site variation alone sometimes does not explain the observed phenotypic variation. The region surrounding the marker on chromosome 14 contains two genes potentially involved in redox homeostasis, a peroxidase (LOC110380531) and a glutaredoxin-3 (LOC110380771), which has not yet been functionally implicated in chlorantraniliprole resistance. Nevertheless, chlorantraniliprole can induce oxidative stress and antioxidant responses in lepidopteran insects (Zhao et al. 2022), and altered oxidative enzyme activity has also been associated with chlorantraniliprole resistance (Wang et al. 2020), which corroborates the possible implication of peroxidase and a glutaredoxin-3 in diamide resistance. Transcriptomic studies further support the involvement of general metabolic responses in resistant *P. xylostella* (Shabbir et al. 2021) and glutathione-dependent systems are well established as components of cellular protection against insecticide-induced oxidative stress (Enayati et al. 2005; Pavlidi et al. 2018). Moreover, other functionally validated non-target-site mechanisms of chlorantraniliprole resistance have been reported (Mallott et al. 2019), corroborating our findings of LOC110380531 and LOC110380771 being associated with this pesticide resistance in *H. armigera*. Together, the associations with chromosomes 14 and 28 support a multilocus architecture in which *RyR*-mediated resistance may coexist with additional non-target-site mechanisms.

The distribution of the resistance-associated *RyR* variants provides further insight into the genetic variation from which R-CHL evolved. Although these variants were detected in only four of the 12 populations screened, their frequencies varied considerably, reaching 0.38 in MT06, the F1 progeny of the field population collected in Itiquira, Mato Grosso, in 2021 that was subsequently used to establish R-CHL (Führ et al. 2026). The same population already showed reduced susceptibility to chlorantraniliprole at F2, with a 7-fold resistance ratio, and responded rapidly and progressively to laboratory selection, reaching 26-fold resistance at F3, 75-fold at F10, and 149-fold at F11 (Führ et al. 2026). Correspondingly, the chlorantraniliprole LD₅₀ increased from 0.157 µg a.i./cm² at F2 to 3.125 µg a.i./cm² at F11, representing an approximately 20-fold increase relative to the initial field-derived population (Führ et al. 2026). The presence of the resistance-associated *RyR* variants at appreciable frequency before selection, together with the rapid phenotypic response observed over subsequent generations, is consistent with chlorantraniliprole selection acting on resistance determinants already segregating within the founding population. However, because we did not track *RyR* allele frequencies across the selection experiment, we cannot establish a direct relationship between increased resistance and enrichment of G4946E/F4947I. Thus, while the data strongly support selection on pre-existing genetic variation, we cannot exclude contributions from additional loci, including those suggested by the genome-wide association analysis.

A different pattern emerged for flubendiamide, indicating that *RyR* coding variation within the region examined here cannot universally explain resistance across diamide insecticides. The Flub-R strain, originally selected from a Brazilian field population collected in 2016, exhibited more than 50,000-fold resistance to flubendiamide despite carrying no amino acid substitutions within the sequenced *RyR* region spanning transmembrane domains II–VI (Abbade-Neto et al. 2022). In contrast, R-CHL, which carries the G4946E/F4947I substitutions and was selected with chlorantraniliprole, exhibited more than 23,000-fold cross-resistance to flubendiamide. These contrasting observations indicate that similar high-level flubendiamide-resistance phenotypes can occur in markedly different *RyR* genetic backgrounds. Some studies in different lepidopteran species show that G4946E is known to reduce *RyR* sensitivity to flubendiamide to a different magnitude than to chlorantraniliprole (Steinbach et al. 2015b; Zuo et al. 2017; Jiang et al. 2021; Wang et al. 2026). This may therefore contribute to the strong cross-resistance observed in R-CHL to flubendiamide when compared to chlorantraniliprole. Furthermore, radioligand binding studies revealed that the G4946E mutation significantly affects both diamide-specific binding and its concentration-dependent allosteric regulation of ryanodine binding (Steinbach et al. 2015b). In contrast, our findings suggest that resistance to the same diamide compound can arise through alternative genetic routes, and that the contribution of target-site variation may depend on the genetic background and resistance history of individual populations.

Consistent with the existence of a new genetic basis for flubendiamide resistance. The strong outlier on chromosome 2 supports a major differentiated region that could contribute to diamide resistance dynamics. Importantly, this association independently recapitulates our previous identification of chromosome 2 as a major resistance-associated region in *H. armigera*, where both controlled genetic crosses and genome-wide analyses implicated this chromosome in the response to flubendiamide (Amado et al. 2025). The recurrence of the same genomic region across distinct populations and complementary experimental designs substantially strengthens the evidence that chromosome 2 harbours one or more determinants contributing to flubendiamide resistance. This interpretation is further supported by the pronounced differentiation observed on chromosome 2 in the genome-wide *F_ST_* scan and GEA analysis, indicating that variation in this region also distinguishes field populations differing in their resistance phenotype against an otherwise weakly differentiated genomic background. Together with the absence of amino-acid substitutions within the examined *RyR* region of the highly resistant Flub-R strain, these independent lines of evidence support a model in which high-level flubendiamide resistance in *H. armigera* can evolve through genetic mechanisms distinct from the canonical *RyR* target-site substitutions (Abbade-Neto et al. 2022; Amado et al. 2025). The causal variant(s) underlying the chromosome 2 signal remain unresolved, but its repeated detection across experimental and field datasets makes this region a major priority for elucidating alternative diamide resistance mechanisms.

The pronounced cross-resistance of R-CHL to flubendiamide further illustrates the complexity of the relationship between *RyR* genotype and resistance phenotype. Although R-CHL was selected with chlorantraniliprole and exhibited approximately 149-fold resistance to the selecting compound, its resistance to flubendiamide exceeded 23,000-fold (Führ et al. 2026), demonstrating substantial cross-resistance between these two diamides. G4946E provides a plausible mechanistic basis for at least part of this response, as functional studies in other lepidopteran species have shown that substitutions at this position can markedly reduce *RyR* sensitivity to both chlorantraniliprole and flubendiamide (Steinbach et al. 2015b). Importantly, however, high-level flubendiamide resistance also evolved in Flub-R in the absence of amino-acid substitutions within the *RyR* region examined here.

Together with the strong and recurrent association of chromosome 2 with flubendiamide resistance, this suggests that laboratory selection of Flub-R may have predominantly enriched an alternative resistance mechanism associated with chromosome 2 rather than *RyR* coding variation. In field populations, however, these alternative routes need not be mutually exclusive. The association of flubendiamide survival with genomic variation outside the *RyR* region, together with evidence that *RyR* substitutions can confer cross-resistance to flubendiamide, suggests that field resistance may arise from multiple mechanisms, including both chromosome 2-associated determinants and target-site variation on chromosome 28. Thus, whereas Flub-R may reflect enrichment of one particularly effective non-*RyR* resistance route under laboratory selection, naturally occurring flubendiamide resistance reflects a more heterogeneous genetic architecture in which distinct resistance mechanisms can occur independently or coexist within populations.

In conclusion, our study reveals that diamide resistance in *H. armigera* can arise through heterogeneous but interconnected genetic routes. For chlorantraniliprole, we identified F4947I as a previously undescribed *RyR* substitution associated with diamide resistance, occurring alongside the well-established resistance mutation G4946E. The presence of this novel G4946E/F4947I combination as standing variation in the field-derived population from which R-CHL evolved, together with its predicted disruption of the diamide-binding pocket and the independent association of the *RyR*-containing chromosome 28 region with chlorantraniliprole survival, provides convergent evidence for a target-site component of resistance. Notably, our structural analyses further suggest that F4947I may modify the effect of G4946E, potentially enhancing disruption of chlorantraniliprole accommodation when both substitutions occur together. In parallel, the recurrent association of chromosome 2 with flubendiamide resistance, together with high-level resistance in Flub-R despite the absence of amino-acid substitutions within the examined *RyR* region, demonstrates that high-level resistance can also evolve through alternative genetic mechanisms. More broadly, our results show that sharing a molecular target does not imply sharing a single evolutionary trajectory to resistance, highlighting the need to consider the genetic architecture of resistance at the level of individual insecticides rather than extrapolating across an entire mode-of-action group.

## Materials and Methods

### Strains

Laboratory strains used in this study were obtained from the Arthropod Resistance Laboratory in Piracicaba, São Paulo, Brazil. The *H. armigera* susceptible strain (SUS) was reared under controlled conditions (25 ± 1 °C, 70 ± 10% relative humidity (RH) and 14:10 h light/dark photoperiod) for more than 100 generations without selection pressure from insecticides or Bt proteins and with no contact with wild individuals. The resistant strain (R-CHL) was selected by F_2_ screening bioassay from a population collected from a soybean crop in Itiquira, Mato Grosso, Brazil, and kept under selection pressure in the diagnostic dose of chlorantraniliprole (0.99 µg a.i.cm^-^²) for 18 generations, showing a resistance ratio of 149-fold in the 11th generation and 141-fold in the 18th generation (Führ et al. 2026). This resistant strain (R-CHL) also showed cross-resistance to other diamide insecticides, such as cyantraniliprole and flubendiamide, with resistance ratios of 422-fold and 23,388-fold, respectively (Führ et al. 2026).

### *RyR* partial sequencing spanning transmembrane domains II-VI

Total RNA was extracted from the thorax of adults of strains SUS and R-CHL. We used Ribozol^™^ RNA Extraction Reagent (VWR Life Science), according to its protocol, for RNA separation, followed by DNase I purification (Thermo Scientific). We quantified the total RNA using a spectrophotometer (Epoch) and assessed its integrity by electrophoresis on a 1.5% agarose gel. We normalised the RNA to 500 ng µL^-1^ and used 1 µg total RNA in a 20 µL reaction for cDNA synthesis with GoScript™ Reverse Transcriptase and oligo(dT) (Promega), following the manufacturer’s instructions.

We manually designed primer pairs from the cDNA sequences of the *RyR* of *H. armigera* (GenBank: KF641862.1 and KJ573634.1), covering transmembrane II to VI (Table S1). Primer pairs were used to amplify fragments overlapping each other and covering a 1553 bp *H. armigera RyR* fragment. The PCR reaction mixture contained 100 ng of cDNA, 500 nM of each primer, 10 mM Tris–HCL, 50 mM KCL (1× PCR buffer; Sinapse), 1.5 mM MgCl₂, 0.2 mM dNTP mix (Sinapse), 1 U of Taq DNA polymerase (Invitrogen; 5 U µL⁻¹), and Milli-Q water to a final volume of 25 µL. We ran the reaction under cycling conditions of 10s at 98 °C, followed by 35 cycles of 98 °C for 3s, 57 °C for 35s, and 72 °C for 15s, and a final extension step at 72 °C for 3 min in a thermal cycler (SimpliAmp, Thermo Fisher). We verified the PCR product by 1% agarose gel electrophoresis, purified it using the EasyPure PCR purification kit (Trans), and subjected it to Sanger sequencing using the forward and reverse primers listed in Table S1. The 1,553-bp partial *H. armigera RyR* sequence obtained by Sanger sequencing was translated into amino acids. The partial *RyR* sequences from *H. armigera* (GenBank accession AHB33498) and the strains susceptible (SUS), chlorantraniliprole-resistant (R-CHL), and flubendiamide-resistant (Flub-R) showed high similarity to the corresponding region of *Plutella xylostella* (Linnaeus, 1758) (Lepidoptera: Plutellidae) *RyR* (GenBank number: AET09964), with 91% pairwise amino acid identity. The *P. xylostella RyR* sequence was used as the reference for residue numbering, following the convention commonly adopted for reporting diamide resistance-associated *RyR* mutations across insect species.

### Structural modelling of *RyR* mutations

Structural modelling was performed using AlphaFold3 (Abramson et al. 2024) to investigate the potential effects of the G4946E and F4947I substitutions on the chlorantraniliprole-binding region of the *H. armigera* ryanodine receptor (*RyR*). A 517-amino-acid region spanning residues of *RyR* encompassing the diamide-binding site was modelled for the susceptible sequence and for sequences carrying the G4946E, F4947I, or combined G4946E/F4947I substitutions. Chlorantraniliprole was provided to AlphaFold3 as a ligand using its SMILES representation.

The cryo-electron microscopy (cryo-EM) *RyR* structure reported by Lin et al. (2024), containing experimentally resolved tetraniliprole within the diamide-binding pocket, was used as a structural benchmark. As a positive control for ligand-pose prediction, we modelled the *RyR* sequence corresponding to the experimentally resolved structure with chlorantraniliprole using AlphaFold3 (Abramson et al. 2024). We compared the predicted position of chlorantraniliprole with the experimentally defined diamide-binding pocket to assess whether the modelling approach reproduced ligand localisation within the expected binding region.

Chlorantraniliprole was subsequently modelled independently with the susceptible *H. armigera RyR* sequence and with the G4946E, F4947I, and G4946E/F4947I variants. For each protein–ligand modelling condition, the highest-ranked prediction according to the AlphaFold3 ranking score was selected for structural comparison. The resulting models were compared with the susceptible *RyR* chlorantraniliprole complex to assess substitution-associated changes in the geometry and accessibility of the diamide-binding pocket and in the predicted position and orientation of chlorantraniliprole relative to residues defining the binding site. Predictions and confidence scores are available in figshare (10.6084/m9.figshare.34069758).

### Field population screening for *RyR* gene mutations

After identifying the RyR mutations in the R-CHL strain, we developed a cleaved amplified polymorphic sequence (CAPS) marker to discriminate between homozygous susceptible, heterozygous and homozygous resistant individuals. The marker was designed to amplify a 208-bp fragment of the RyR gene encompassing the identified substitutions, using the primers Chlo-F (5′-AATCAACATAGACTGGCGCTAC-3′) and Chlo-R (5′-TACCTGTTTCCCATTATGCGTG- 3′). Primer specificity was assessed in silico and confirmed by Sanger sequencing of the amplified fragments from SUS and R-CHL individuals. Restriction enzyme screening was subsequently performed in silico using Benchling (Anon 2022), and TaqI (10 U µL⁻¹; Thermo Fisher Scientific, catalogue no. ER0671) was selected based on its ability to discriminate between susceptible and resistant alleles. The CAPS marker was initially validated using laboratory strains and subsequently applied to 221 individuals from 12 field-derived populations collected between the 2015 and 2021 crop seasons.

To investigate the relationship between RyR resistance-associated allele frequencies and phenotypic susceptibility, we integrated previously published and newly generated chlorantraniliprole susceptibility-monitoring data. The diagnostic concentration of 0.99 µg a.i. cm⁻² was established by Pereira et al. (2020), who also conducted and reported susceptibility-monitoring bioassays for field populations collected during the 2014-2018 crop seasons. Susceptibility data for populations collected during the 2019-2021 crop seasons are reported here for the first time. These additional populations were evaluated using the same diagnostic concentration and bioassay protocol established by Pereira et al. (2020), ensuring methodological consistency across the monitoring period. Accordingly, the population-level survival percentages used in the present study were obtained from Pereira et al. (2020) for the earlier crop seasons and from the newly conducted susceptibility-monitoring bioassays for the subsequent crop seasons.

We then examined the relationships between resistance-associated allele frequency, population survival percentage and crop season using correlation analyses implemented with the cor and cor.test functions in R (R Core Team 2025). To further evaluate the ability of the CAPS marker to discriminate between populations with different susceptibility phenotypes, we performed receiver operating characteristic (ROC) analysis using resistance-associated allele frequency as the predictor variable. Population survival percentages at the diagnostic concentration of 0.99 µg a.i. cm⁻² were dichotomised into two categories: populations with survival ≥20% were classified as survival (1), whereas those with survival <20% were classified as dead (0). ROC curves were generated using the roc function from the pROC package (Robin et al. 2011), and the area under the curve (AUC) was calculated using auc. Bootstrap confidence intervals were estimated using ci.auc with 10,000 replicates, and statistical significance was assessed using the pnorm function.

### F_ST_ scan across field populations

We performed an F_ST_ scan using 189 individuals (ENA: ERP162059) (North et al. 2025). Population-level survival percentages were obtained from chlorantraniliprole and flubendiamide susceptibility-monitoring bioassays conducted using the diagnostic concentrations established by Pereira et al. (2020). Survival data for populations collected during the 2014-2018 crop seasons were previously published by Pereira et al. (2020), whereas those for the 2019-2021 crop seasons are reported here for the first time. Based on these phenotypic data, individuals were assigned to two groups according to the survival percentage of their source population. For chlorantraniliprole, individuals originating from populations with survival rates of 1-10% were assigned to Population A, whereas those from populations with survival rates ≥11% were assigned to Population B. For flubendiamide, Population A comprised individuals from populations with survival rates of 1-14%, whereas Population B comprised individuals from populations with survival rates ≥20% (Table S2). SNPs were filtered to retain variants with a minor allele frequency (MAF) ≥0.05 and genotype data available for at least 80% of individuals (--max-missing 0.80).

We used the Weir and Cockerham F_ST_ estimator to estimate genome-wide genetic differentiation, implemented in VCFtools v0.1.16 (Danecek et al. 2011). We calculated pairwise F_ST_ using a sliding-window approach with 50-kb windows and a 10-kb step size to identify highly differentiated genomic regions. To assess the robustness of candidate signals and improve mapping resolution, we repeated the sliding-window analysis using 10-kb windows with a 4-kb step size. We considered genomic regions with consistently elevated F_ST_ values across both window sizes as robust candidates for population-level genetic differentiation.

### Genotype-environment association with field populations across the 2014-2021 crop seasons

We performed genotype-environment association (GEA) analyses using BayPass v2.4 (Gautier 2015) to identify SNPs whose allele frequencies were associated with variation in insecticide survival among field populations, while accounting for population covariance. The analysis included 189 individuals from 28 field populations collected across the 2014-2021 crop seasons (Table S2). The VCF file was filtered using the same criteria described above for the *F_ST_* analysis. After filtering, biallelic SNPs were normalised to the reference genome using BCFtools (Danecek et al. 2021), and genotypes were converted to reference and alternative allele counts for each population. A minimum within-population genotype call rate of 70% was required for a population to contribute allele counts at a given SNP.

Population-level survival percentages were obtained from susceptibility-monitoring bioassays conducted using the diagnostic concentrations established by Pereira et al. (2020): 0.99 µg a.i. cm⁻² for chlorantraniliprole and 2.64 µg a.i. cm⁻² for flubendiamide. Survival data for populations collected during the 2014-2018 crop seasons were previously reported by Pereira et al. (2020), whereas data for the 2019-2021 crop seasons are presented here for the first time. The same diagnostic concentrations and bioassay protocol were maintained throughout the monitoring period to ensure methodological consistency. These population-level survival percentages were used as phenotypic covariates in the GEA analyses and transformed to the logit scale, log[*p*/(1 − *p*)], where *p* represents the proportion of individuals surviving insecticide exposure within each population. To account for covariance in allele frequencies among populations resulting from their shared demographic history, we estimated the population covariance matrix (Ω) from a subset of approximately independent SNPs distributed across the genome. SNPs were pruned for linkage disequilibrium using PLINK2 (Chang et al. 2015) with a 50-SNP sliding window, a five-SNP step size, and an *r^2^* threshold of 0.20. Up to 75,000 SNPs were retained for covariance estimation. When LD pruning did not provide adequate genome-wide representation, we used systematic thinning to obtain markers distributed across the genome.

The Ω matrix was estimated from three independent BayPass core-model runs using 10,000 burn-in iterations, a thinning interval of 25, and 1,000 sampling iterations. The final covariance matrix was obtained by averaging the Ω matrices estimated across the three independent runs. Associations between allele-frequency variation and chlorantraniliprole and flubendiamide survival were subsequently evaluated using the BayPass importance sampling (IS) model, incorporating the estimated Ω matrix to account for population structure. To accommodate the genome-wide dataset, SNPs were analysed in blocks of up to 75,000 markers. We performed three independent IS analyses and retained the median association statistic across the three runs for each SNP to reduce variation attributable to MCMC sampling. Evidence for association was evaluated using the Bayes factor expressed in decibans [BF(dB)], with 20 dB corresponding to 100:1 odds (Wod 1985; Gautier 2015). We considered SNPs with BF ≥ 20 dB as preliminary candidates associated with variation in the survival of chlorantraniliprole and flubendiamide.

## Author contribution

**Douglas Amado:** Writing – review & editing, Writing - original draft, Visualisation, Validation, Supervision, Software, Resources, Project administration, Methodology, Investigation, Formal analysis, Data curation, Conceptualisation. **Anne Caroline Santos Küll:** Validation, Methodology, Investigation, Data curation. **Fábio Miguel Führ:** Writing - review & editing, Formal analysis, Conceptualisation. **Victor Flores:** Methodology, Formal analysis, Visualisation. **Erika Pinheiro de Castro:** Formal analysis, Writing – review & editing, Conceptualisation. **Alberto Soares Corrêa:** Formal analysis, Funding acquisition, Project administration, Supervision, Writing – review & editing, Conceptualisation. **Chris D. Jiggins:** Formal analysis, Funding acquisition, Project administration, Supervision, Writing – review & editing, Conceptualisation. **Celso Omoto:** Writing - review & editing, Writing - original draft, Validation, Supervision, Project administration, Funding acquisition, Formal analysis, Data curation, Conceptualisation.

## Conflicts of interest

The authors declare no competing interests.

## Funding statement

This research was financed by the Biotechnology and Biological Sciences Research Council (BBSRC: BB/V001329/1) and the São Paulo Research Foundation (FAPESP: 2020/00708-7) and (FAPESP: 2019/18282-9). The National Council for Scientific and Technological Development (CNPq) offered a Doctoral Scholarship to DA (142590/2019-3), and Research Fellowships to CO (308459/2025-3) and ASC (307115/2026-7). Bilateral BBSRC-FAPESP: Population genomic monitoring of soybean pests (UKRI2955). VF was funded by a Gordon and Betty Moore Foundation Investigator Award to Ross F. Waller (https://doi.org/10.37807/GBMF9194).

## Acknowledgements

We thank the Brazilian Insecticide Resistance Action Committee (IRAC-BR) for providing *Helicoverpa armigera* populations for this research.

## Notes

https://figshare.com/articles/online_resource/AlphaFold3_modelling_of_RYR_with_chlorantraniliprole/34069758

### Competing Interest Statement

The authors have declared no competing interest.

### Summary of Updates

Diamide insecticides target insect ryanodine receptors (RyRs) and are widely used against lepidopteran pests, yet it remains unclear whether compounds sharing this molecular target evolve resistance through the same genetic routes. Here, we integrated RyR sequencing, structural modelling and population genomics to investigate resistance to chlorantraniliprole and flubendiamide in the global agricultural pest *Helicoverpa armigera*. We identified G4946E together with F4947I, a previously undescribed resistance-associated RyR substitution, in a chlorantraniliprole-resistant strain. Structural modelling indicated that F4947I may modify G4946E's effects on the diamide-binding pocket, while field and population-genomic evidence independently implicated the RyR-containing chromosome 28 region in chlorantraniliprole resistance. By contrast, high-level flubendiamide resistance occurred without amino acid substitutions in the RyR region examined, and field survival was predominantly associated with chromosome 2, independently recapitulating a previously identified resistance locus. Together, our findings reveal that resistance to insecticides sharing the same molecular target can evolve through distinct but potentially overlapping genetic routes, highlighting the limitations of predicting cross-resistance from target-site variation alone and the need for broader genomic approaches to resistance surveillance.

