## Supplemental Table 1 for "A shared molecular target but distinct genetic routes to diamide resistance in *Helicoverpa armigera* (Lepidoptera: Noctuidae)"

### Supplementary material

**Table S1.** List of primers used in Sanger sequencing.

| Primer | Sequence 5'- 3' | Amplification length (pb) |
|---|---|---|
| HA1-F | CTCAAGAGAGGATGGAAGC | 728 |
| HA1-R | AGAAGGAAGTCTTCTCCATGC | |
| HA2-F | TCGCTCGTAAGCTGGAAT | 756 |
| HA2-R | AGACCCTGGAGAATAGCG | |
| HA3-F | TTCCGCAAGTTCTACGTGC | 510 |
| HA3-R | TTCGGAAACAATCTCCCACG | |

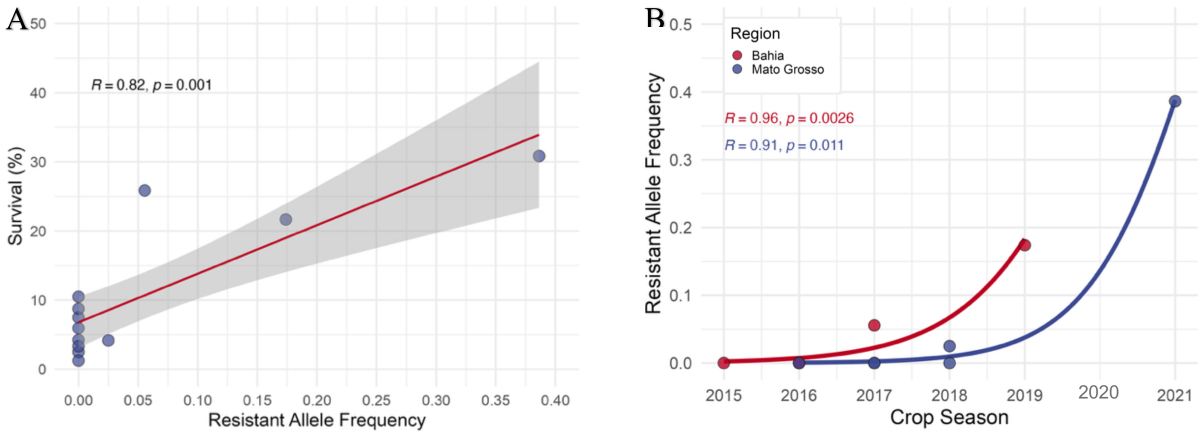


**Figure S1.** Resistant allele frequency predicts survival and increases over time in field populations. A) Relationship between resistant-allele frequency and survival (%) of field populations exposed to a diagnostic dose of chlorantraniliprole. Points represent field populations. The solid line shows the fitted linear regression, and the shaded band indicates the 95% confidence interval. Pearson's correlation coefficient is reported. B) Temporal dynamics of resistant-allele frequency across crop seasons (2015–2021). Points represent seasonal allele-frequency estimates. The solid line shows the fitted linear regression, and the shaded band indicates the 95% confidence interval. Pearson's correlation coefficient quantifies the association between allele frequency and time.

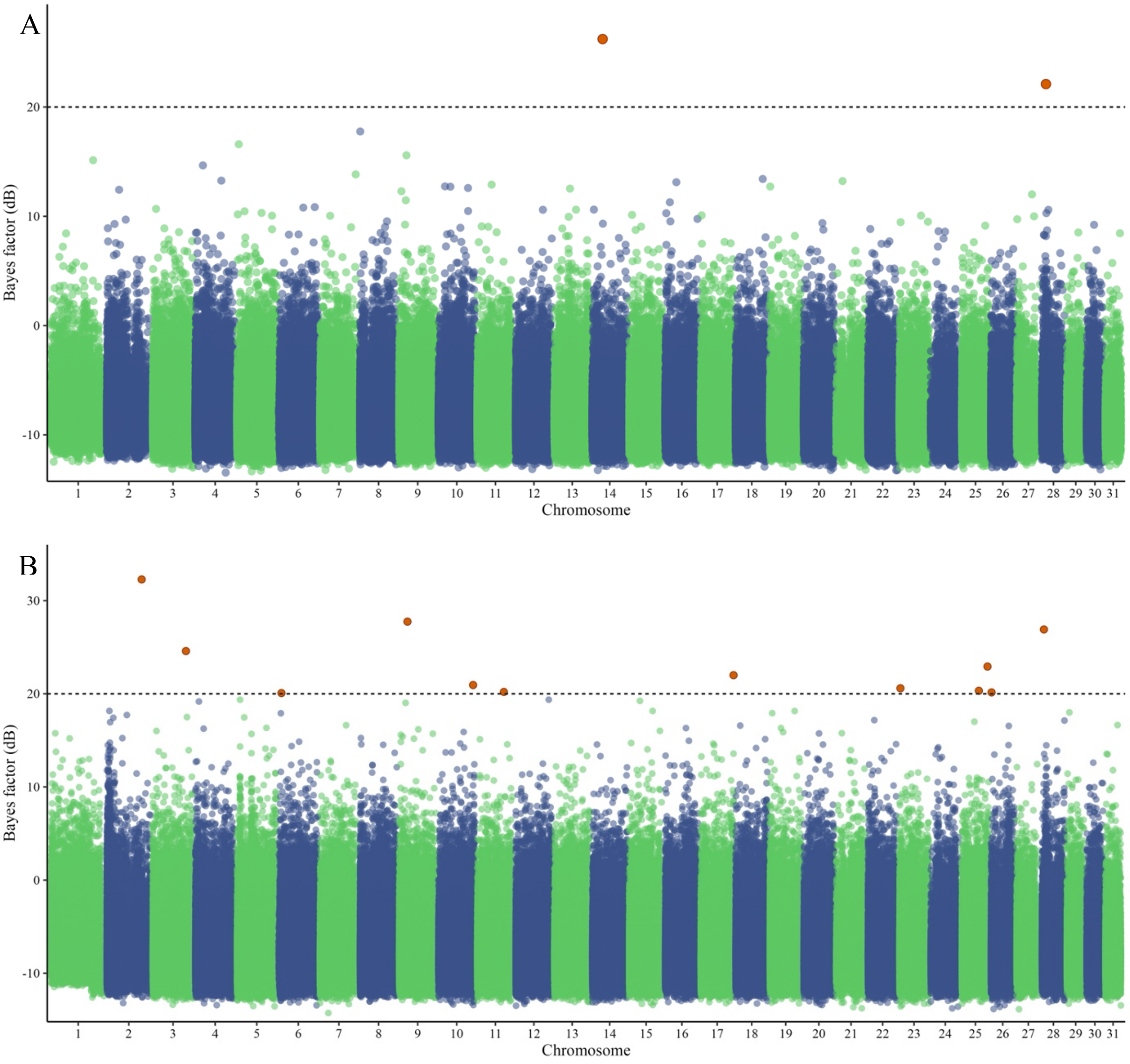


**Figure S3.** Genome-environment associations between allele frequencies and diamide insecticide survival in field populations of *H. armigera*. Manhattan plots show BayPass genotype–phenotype associations with population-level survival to (A) chlorantraniliprole and (B) flubendiamide. Each point represents a SNP, with chromosomes on the x-axis and Bayes factors [BF(dB)] on the y-axis. The horizontal dashed line indicates the association threshold of BF = 20 dB, and SNPs exceeding this threshold are highlighted in orange.
